# Specificity Principles in RNA-Guided Targeting

**DOI:** 10.1101/055954

**Authors:** Namita Bisaria, Inga Jarmoskaite, Daniel Herschlag

## Abstract

RNA-guided nucleases (RGNs) provide sequence-specific gene regulation through base-pairing interactions between a small RNA guide and target RNA or DNA. RGN systems, which include CRISPR-Cas9 and RNA interference (RNAi), hold tremendous promise as programmable tools for engineering and therapeutic purposes. However, pervasive targeting of sequences that closely resemble the intended target has remained a major challenge, limiting the reliability and interpretation of RGN activity and the range of possible applications. Efforts to reduce off-target activity and enhance RGN specificity have led to a collection of empirically derived rules, which often paradoxically include *decreased* binding affinity of the RNA-guided nuclease to its target. Here we demonstrate that simple kinetic considerations of the targeting reaction can explain these and other literature observations. The kinetic models described provide a foundation for understanding RGN systems and a necessary physical and functional framework for their rational engineering.

## INTRODUCTION

Over the past decades, RNA-guided nucleases (RGNs) have emerged as powerful and versatile tools for genome editing and for manipulation of gene expression. RNAi, CRISPR, and ribozyme targeting utilize short oligonucleotides to guide enzymatic activity to specific DNA and RNA sequences through base-pairing interactions. The established rules for Watson-Crick base-pairing and their predictable energetics have led to the prevailing perspective that RGNs and related systems are readily programmable and inherently specific. Nevertheless, experimental studies have exposed limitations to RGN specificity, revealing persistent targeting of unintended sequences that resemble the target RNA or DNA sequence (‘off-targets’) (Hu et al., 2016; Jackson et al., 2003). This limited specificity is one of several key challenges that must be overcome to allow RGN-based technologies to achieve their full impact in the laboratory, in engineering, and ultimately as therapies (Kleinstiver et al., 2016; Slaymaker et al., 2016; Yin et al., 2016).

A series of successes in reducing off-target cleavage have been reported, most recently in the CRISPR-Cas9 field using changes in the protein, RNA guide, or both (Doench et al., 2016; Fu et al., 2014; Guilinger et al., 2014; Hu et al., 2016; Kleinstiver et al., 2016; Mali et al., 2013; Ran et al., 2013; Slaymaker et al., 2016). A common theme that emerged from studies of CRISPR-Cas9 targeting is that specificity of targeting increases upon *weakening* interactions of the Cas9-RNA complex with the target DNA— e.g., by using truncated RNA guides (Fu et al., 2014; Guilinger et al., 2014; Herschlag, 1991; Miller et al., 2003; Pattanayak et al., 2011; Pedersen et al., 2014; Østergaard et al., 2013) or by introducing mutations to the protein-DNA interface (Kleinstiver et al., 2016; Slaymaker et al., 2016; Yin et al., 2016). This counterintuitive observation has been rationalized in terms of an “excess energy” model, postulating that binding energy beyond a certain threshold stabilizes binding to both correct and incorrect targets (Doench et al., 2016; Fu et al., 2014; Guilinger et al., 2014; Hu et al., 2016; Kleinstiver et al., 2016; Mali et al., 2013; Ran et al., 2013; Slaymaker et al., 2016). Similar observations have previously been made for RNAi, RNase H, and ribozyme targeting, and in the related fields of TALEN and zinc-finger-nuclease targeting (Fu et al., 2014; Guilinger et al., 2014; Herschlag, 1991; Miller et al., 2003; Pattanayak et al., 2011; Pedersen et al., 2014; Østergaard et al., 2013). However, a physical model that explains the “excess energy” phenomenon in CRISPR-Cas9 and RNAi has not been presented. More generally, a mechanistic explanation that takes into account the kinetics of individual steps of the targeting process will help transform isolated successes into generalizable approaches that will in turn direct the development of new rational strategies for improving the specificity of RGNs.

Here, we demonstrate the importance—and power—of kinetic considerations for determining and improving RGN specificity. This work builds on prior discussions of specificity for ribozyme targeting (Herschlag, 1991). We first discuss the inadequacy of binding thermodynamics alone to account for specificity of RGN targeting, and we then describe a simple kinetic model that includes steps in addition to binding and is supported by published biochemical studies of RGNs. We explain recent improvements in CRISPR-Cas9 and RNAi specificity in the context of this kinetic model and discuss possible biological implications of differences in the targeting kinetics—and consequently, specificities—of different RGNs. The models described here provide a conceptual framework for future engineering to enhance the specificity of RGN systems and for accurate predictions of RGN targets in vivo.

## RESULTS AND DISCUSSION

### Thermodynamic and kinetic factors in targeting specificity

An intuitive approach to understanding and predicting RGN specificity—and one that is generally taken, explicitly or implicitly, considers the thermodynamics of target versus off-target recognition. The target binding step is depicted for a generic RGN system in Figure 1A.

**Figure 1.**
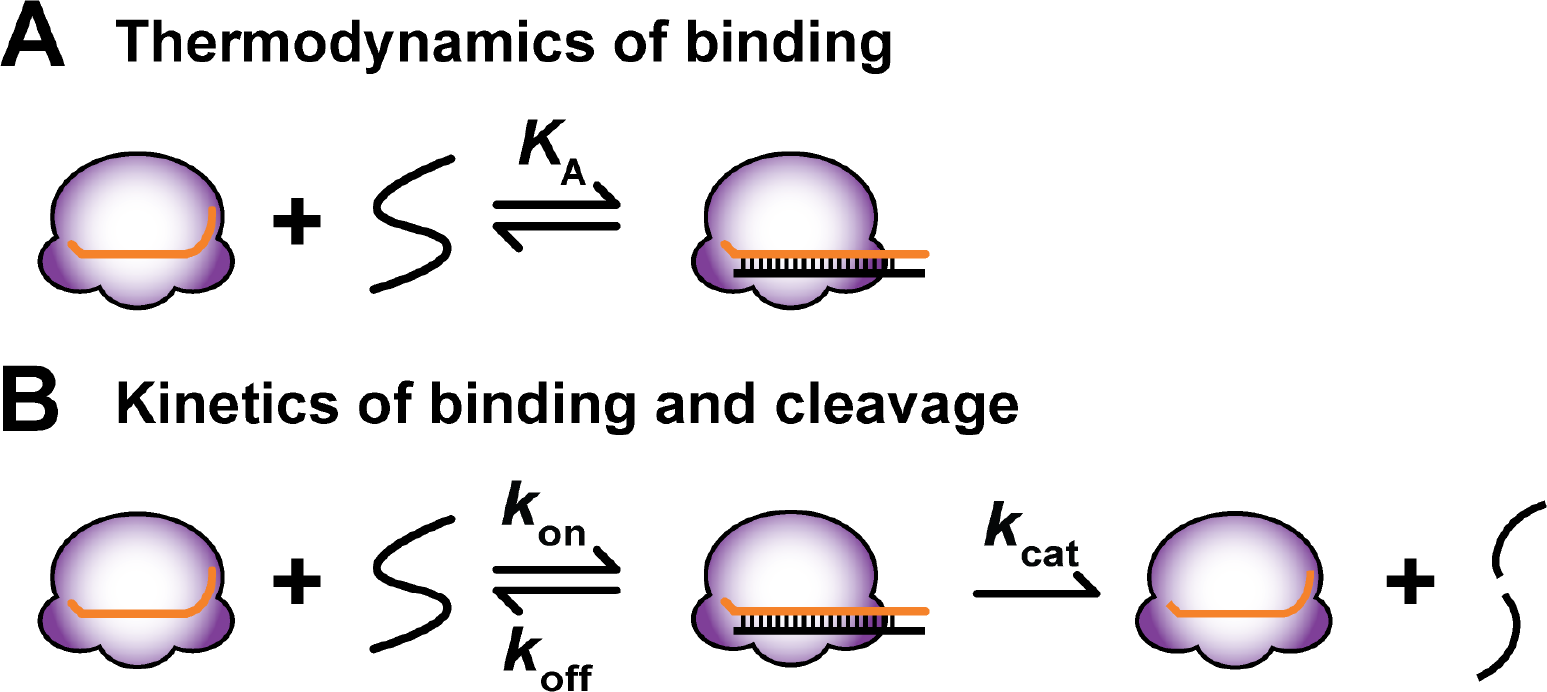
RGN targeting from thermodynamic (**A**) and kinetic (**B**) perspectives. Ka: equilibrium association constant for recognition of a target (black) by a generic RGN (purple), loaded with an RNA guide (orange). *k*_on_, *k*_off_, *k*_cat_: rate constants for target binding, dissociation and cleavage, respectively.

If the thermodynamics of target recognition were the sole determinant of specificity, specificity for the correct versus off-target site would increase with the number of residues involved in recognition as the number of mismatches encountered between the guide and off-target sites would increase. Figure 2A uses this thermodynamic model and shows predicted target affinity by guide RNAs of varying lengths, plotting the binding affinity for a matched target and three off-target nucleic acids with one, two, or three mismatches. Consistent with simple thermodynamic intuition, the binding affinity for the target increases steadily as the number of base pairs utilized in targeting increases (Figure 2A; black dots). Binding for the mismatched targets (colored dots) also increases with guide RNA length, but not monotonically, as the increase in binding energy is interrupted when a mismatch is encountered. Thus, binding specificity, or the difference in binding affinity between the matched and mismatched targets, increases as the guide RNA is lengthened and as the probability of encountering additional mismatches increases (Figure 2B). According to the thermodynamic perspective, specificity can be 'dialed’ to be as high as desired against off-target competitors by continuing to lengthen the guide, which allows the incorporation of more and more mismatches against off-targets. However, as emphasized in the Introduction, there is empirical evidence that this pattern is not followed and that *shortening* recognition sequences or weakening binding can instead increase apparent target specificity.

**Figure 2.**
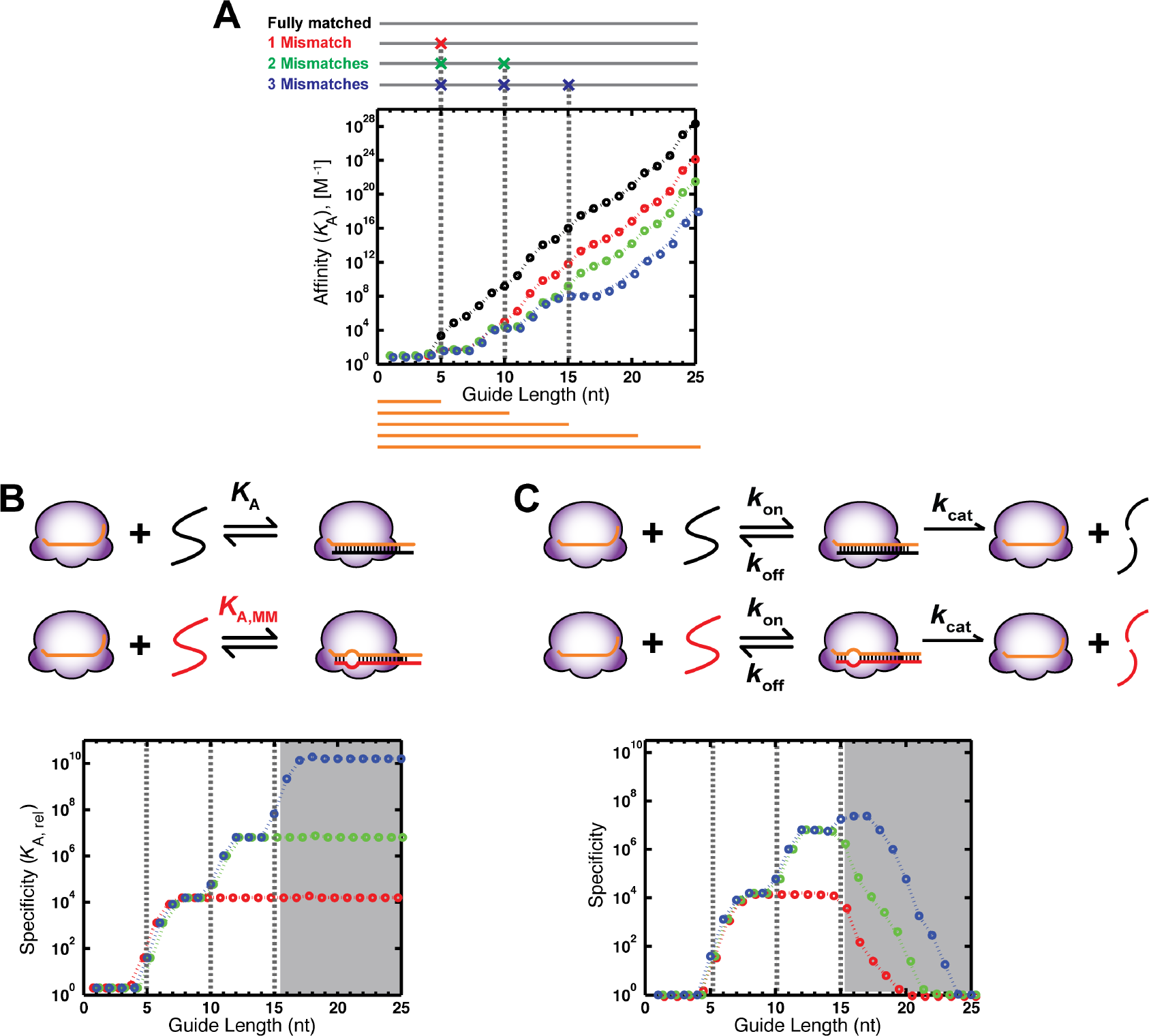
Thermodynamic and kinetic perspectives of RGN specificity. **A**. Model for changes in the binding affinity for target and off-target sites with increasing length of the RNA guide. Orange lines show guides of representative lengths. The targets (top) include one that is fully complementary (black) to the guide RNA, and off-target sites (colors) that contain one, two, or three mismatches at positions 5, 10 and 15 (denoted with gray dashed lines). Binding affinities (*K*_A_) were calculated for RNA-RNA helices using DinaMelt (Markham and Zuker, 2005). The sequence of let-7a miRNA was used and was extended at the 3’ end (<u>UGAGGUAGUAGGUUGUAUAGU</u>*AGCA*) to simulate affinities for duplex lengths greater than 21 bp. **B**. Thermodynamic specificity model. Scheme of binding of an RGN to a matched (black) and singly mismatched (red) target. Binding specificity for the fully complementary target versus each mismatched target was calculated with values from panel (A) (Specificity = *K*_A, rel_ = *K*_A_(matched)/*K*_A_(mismatched)). Color scheme is the same as in (A). The gray area indicates guide lengths that display differences in specificity between the thermodynamic and kinetic models (panel (C)). **C**. Kinetic specificity model. Kinetic scheme of binding followed by a cleavage step. Here, specificity is the ratio of *k*_cat_/*K*_M_ values for the fully complementary target relative to each mismatched target (see text). The simulation assumes constant *k*on and *k*cat values, while *k*off changes with the number of base-pairs and mismatches, as predicted from thermodynamic stabilities in (B) (*k*_off_ = *k*_on_/*K*_A_). For the purpose of illustrating general concepts of specificity, we assume that mismatches do not affect the cleavage rate and simulate all binding affinities based on nearest-neighbor predictions for RNA-RNA base-pairing. In reality, these target affinities are influenced by the RGN protein, the type of nucleic acid, and these effects influence specificity, as discussed later in the text. For calculations and rate constants used in simulations, see Experimental Procedures; for simulations with different *k*on and *k*cat values, see Figure S1.

RGN systems involve additional steps that follow binding such as cleavage, miRNA-dependent repression and degradation (Carthew and Sontheimer, 2009), and engineered activities of cleavage-inactivated CRISPR-Cas9 systems (Dominguez et al., 2016). For simplicity, we focus on RGN-catalyzed cleavage (Figure 1B), but emphasize that the concepts apply to any targeting system with an irreversible step. Figure 2C depicts the specificity that results when this additional step is taken into account, by including in the model a cleavage step and its corresponding rate constant in addition to rate constants for the binding step (with the off-rates derived from a constant on-rate and the binding affinities from Figure 2A; see Experimental Procedures). The resulting specificity profile (Figure 2C) is very different than that based solely on thermodynamic considerations (Figure 2B). For shorter guides, specificity rises with increasing guide length, as expected from the binding considerations in Figure 2A and is indeed identical to that observed in Figure 2B; however, as guide length is increased further, specificity decreases rather than continuing to increase or leveling off (Figure 2C, grey background). While one may describe this problem as arising from “excess binding energy”, this term only describes the observation and does not provide a physical model to account for the behavior. In the next section, we describe the kinetic principles that underlie the relationship between binding energy and specificity in terms of a simple kinetic model that can be used to guide and interpret future experiments.

### Kinetic model of RGN specificity

We would like an RGN, such as that depicted generically in Figure 3A, to discriminate between a fully matched, intended target and the large number of potential off-targets that contain mismatches. Two kinetic regimes exist, which we describe indepth below, and the extent to which an RGN will be able to discriminate between the intended target and potential mismatched off-targets will depend on the kinetic regime that the RGN is in.

**Figure 3.**
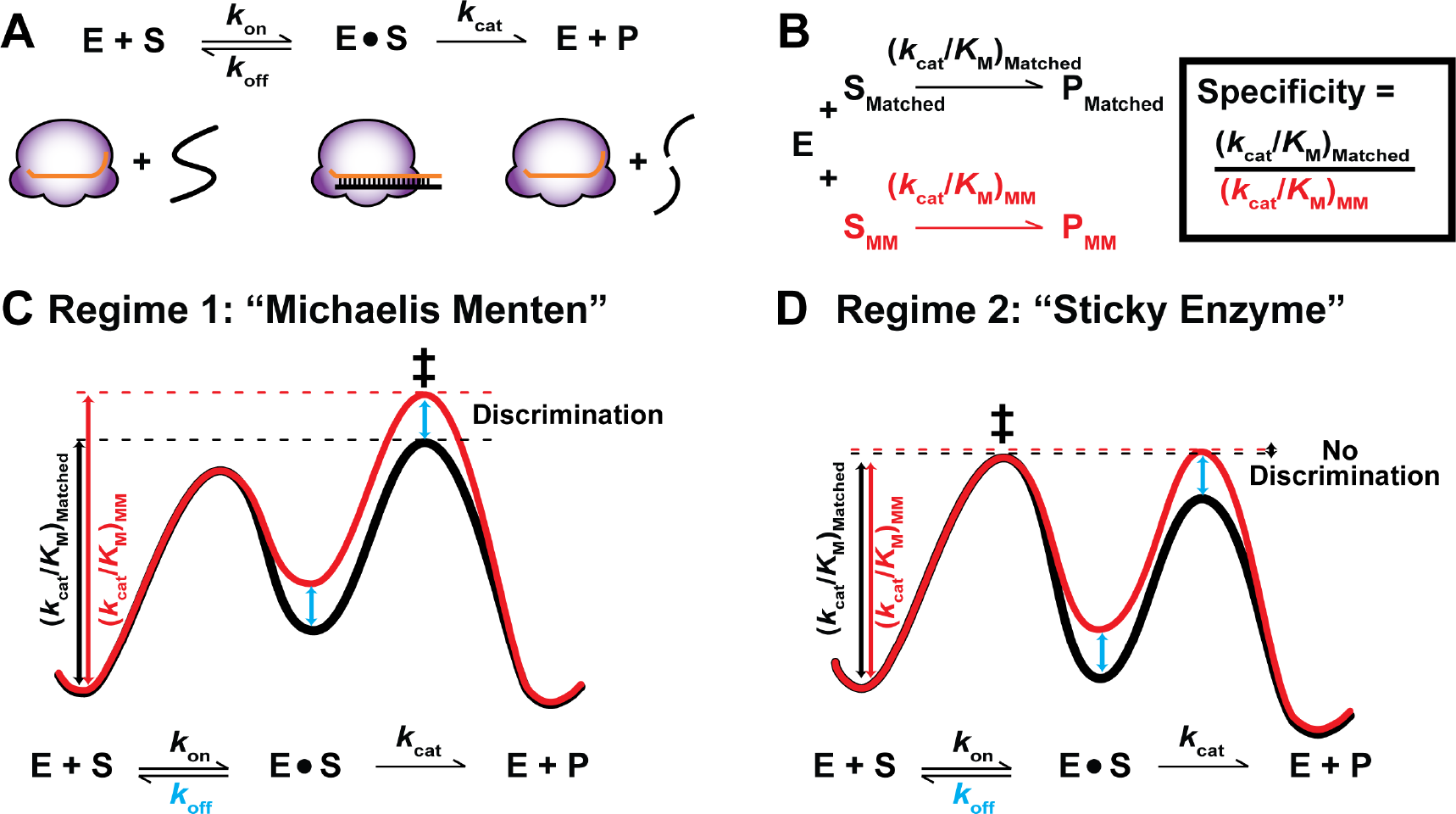
Two kinetic regimes and their implications for specificity. **A**. Overall kinetic scheme for binding and cleavage. **B**. Specificity is defined as the ratio of *k*_cat_/*K*_M_ values for a matched (black) versus mismatched (MM, red) target (Fersht, 1999). **C**. Free energy diagram for a matched versus mismatched target following Michaelis-Menten kinetics (Regime 1). *k*_off_ is larger than *k*_cat_ (for both S_Matched_ and S_Mm_), allowing for binding equilibration prior to cleavage. Overall discrimination (difference between the black and red vertical arrows, indicating *k*_c_JK_u_ values) corresponds to the difference in thermodynamics at the binding step (blue arrow). **D**. Free energy diagram for a matched versus mismatched target following Regime 2 kinetics. In this regime, *k*_off_ is smaller than kcat, so that binding equilibration cannot occur prior to cleavage and specificity is lost. For simplicity, in Panels (C) and (D) we assume that mismatches primarily affect the rate of dissociation (*k*_off_) of the RGN from the target (i.e., a reduced height of the barrier between E•S and E + S for the mismatched relative to the matched target) and use identical *k*_on_ and *k*_cat_ values for the matched and mismatched targets (i.e., identical heights of the barriers between E + S and E•S (*k*_on_) and between E•S and E + P (*k*_cat_)). This assumption for *k*_cat_ appears to hold for mismatches away from the cleavage site in the Argonaute RGN (Wee et al., 2012), and possiblevariations in *k*_on_ are addressed below. The energetic barrier for *k*_cat_/*K*_M_ is designated with vertical lines from the ground state (E + S) to the transition state (‡) of the reaction.

Discrimination, or specificity, for a process is determined by the ratio of ‘*k*_cat_/*K*_M_’ values for the reaction with two competing substrates—in this case, a correct, or matched (S_Matched_) versus incorrect, or mismatched target (S_MM_) (Figure 3B; (Fersht, 1999)). The *k_cat_/K_M_* value represents the overall rate for a reaction and takes into account an enzyme's ability to bind, dissociate and, in this case, cleave a target, in terms of rate constants for each of these processes (*k*_on_, *k*_off_, and *k*_cat_, respectively). For a simple presentation of the concepts, we assume that *k*_cat_ is the same as the rate of cleavage. Below, we illustrate how the relative magnitudes of the rate constants that comprise *k*_cat_/*K*_M_ determine the extent of discrimination between matched and mismatched targets. These models can be expanded to take into account more complex scenarios, including those in which additional steps such as product release contribute to k_cat_, which defines the maximum turnover rate constant (Haley and Zamore, 2004; Salomon et al., 2015; Sternberg et al., 2014; Wee et al., 2012).

Two kinetic regimes can be defined based on the relative magnitudes of the rate constants *k*_off_ and *k*_cat_: one that allows differences in binding affinities between a correct and incorrect target to manifest in specificity and one that masks these differences, thereby reducing or eliminating discrimination. The kinetic regime that allows for discrimination was originally described by Michaelis and Menten (Michaelis et al., 2011). Figure 3C illustrates Michaelis-Menten kinetics for an RGN reaction using a free energy-reaction diagram, where the wells represent stable states and the peaks represent the barriers for transitions between those states. The heights of individual barriers are inversely proportional to the log of the rate constant for the corresponding transition, and *k*_cat_/*K*_M_ (the overall efficiency of catalyzing the reaction of a particular substrate) is represented by the total height of the free energy barrier in going from the unbound state (E + S) to the highest reaction barrier, or transition state (‡), in the reaction.

The RGN (‘E’) can bind either the matched (black curve) or the mismatched (red curve) substrate. Once the bound complex is formed, there are peaks on either side of the ‘E•S’ complex, one for dissociation to E and S (*k*_off_) and the other for cleavage to E + P (*k*_cat_). The central feature of Michaelis-Menten kinetics is that the peak for cleavage is higher compared to dissociation—both for the matched and the mismatched target. This means that the bound E•S_Matched_ and E•S_MM_ complexes dissociate faster than they are cleaved (have smaller barriers for dissociation than cleavage) and that cleavage is the rate-limiting step. Stated another way, the free and bound state of E and S can bind and dissociate many times prior to cleavage and so are in equilibrium. Since E (our RGN) can equilibrate its binding to S_Matched_ and S_MM_ before it cleaves, there is a preference for cleaving S_Matched_ over S_MM_ that matches the thermodynamic preference for binding S_Matched_ over S_MM_. This scenario corresponds to the regions in Figure 2B and 2C with shorter and thus less stably base-paired RNA guides (white backgrounds), where the thermodynamic (i.e., binding) specificity (Figure 2B) is the same as the reaction specificity (Figure 2C).

The reaction properties that underlie the second kinetic regime, in which reaction specificity diverges from binding specificity, are depicted in Figure 3D. This scenario will occur with a longer guide sequence or with stronger interactions with a given guide (gray regions in Figures 2B and 2C). In this regime, once the E•S_Matched_ or E•S_MM_ complex is formed, both are preferentially cleaved—i.e., the barriers to return to the free E + S_Matched_ or E + S_MM_ states are higher than the barrier for cleavage to form E + P_M_ or P_MM_. In other words, once either S_Matched_ or S_MM_ bind, they are ‘stuck’ to the enzyme and are cleaved instead of dissociating, essentially every time they bind E. Such enzymes are sometimes referred to as ‘sticky enzymes’ (and the kinetic regime is referred to sometimes as “Briggs-Haldane” kinetics (Briggs and Haldane, 1925)).

So, the ‘excess binding energy’ that was previously noted can correspond to scenarios where binding is so strong that both the matched and mismatched nucleic acids are ‘sticky’ and cleaved at their binding rate (i.e., each time they bind). Indeed, published results provide evidence for both Regime 1 and Regime 2 kinetics in different RGNs, and the kinetic regime for different RNAi systems may have been tuned through evolution to match functional demands for speed versus specificity. We also note that more complex scenarios are possible. E.g., mismatches can lower binding rate constants (*k*_on_) and can slow cleavage (*k*_cat_), thereby giving some specificity. These and other more complex scenarios can be described in terms of reaction profiles like those in Figure 3C and 3D to analogously describe and predict specificities.

### Kinetics and specificity of Argonaute proteins

The most extensive RGN biochemical studies to date have focused on the target binding and cleavage activities of Argonaute enzymes. A comparative study of fly Ago2 and mouse AGO2 enzymes by Wee *et al.*, provides real-world examples of the two kinetic regimes discussed above and the kinetics of each enzyme are suggestive of their in vivo roles in each organism (Wee et al., 2012). In vitro, fly Ago2 cleaves a fully complementary target at a rate that is almost three orders of magnitude faster than dissociation (Figure 4A, *k*_cat_ versus *k*_off_). Thus, fly Ago2 is a “sticky” enzyme: for every thousand fully complementary targets that bind, ~999 are cleaved and only one is expected to dissociate (Wee et al., 2012). The difference between cleavage (*k*_cat_) and dissociation (*k*_off_) is so large that even for targets with multiple mismatches, cleavage can occur before the target has an opportunity to dissociate.

**Figure 4.**
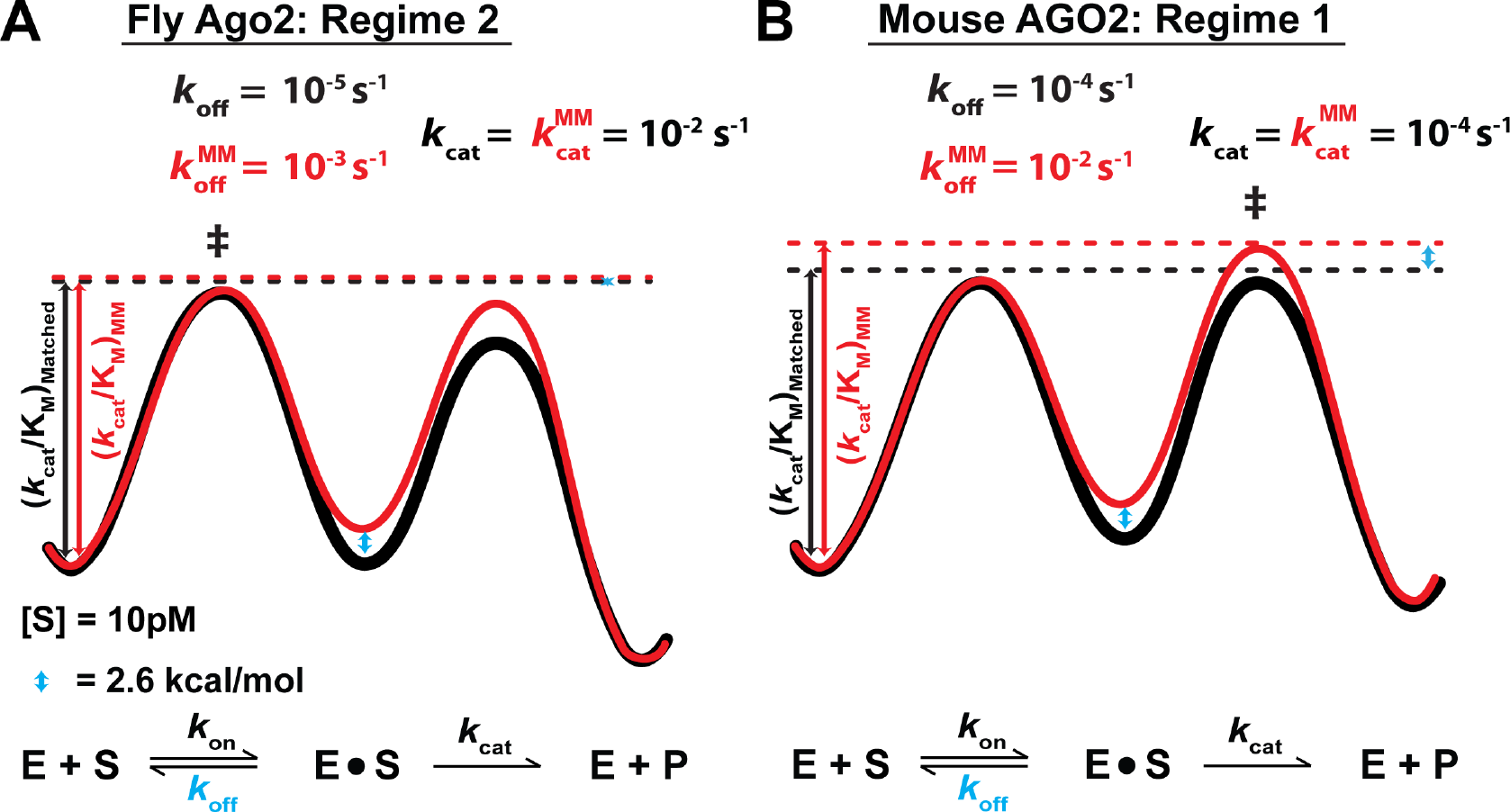
Kinetic regimes of Argonaute proteins from fly and mouse. **A**. Free energy diagram for the silencing reaction by Fly Ago2. Profiles for binding and cleavage of fully matched target (black) and a target containing a mismatch (red) are shown, assuming that the mismatch increases the dissociation rate by 100-fold (likely an overestimate but useful for visualizing the changes). Rate constants for all steps of the fully matched target are taken from Wee et al. (Table S1) (Wee et al., 2012), assuming a RISC concentration of 10 pM (corresponding to approximate concentration of a lowly expressed miRNA) (Wee et al., 2012). We assume that turnover rate (*k*_cat_) is unaffected by this mismatch as *k*_cat_ is generally not changed by single mismatches outside the central region of the guide RNA (guide positions 9-11) (Ameres et al., 2007; Haley and Zamore, 2004; Wee et al., 2012). **B**. Free energy diagram for miRNA-dependent binding and cleavage by mouse AGO2. Source and assumptions as in (A).

On the surface, this absence of high specificity seems troubling, as we typically associate specificity with biological processes. However, fly Ago2 has been implicated in siRNA-mediated silencing for defense against viral infection, whereby foreign RNA is processed by the RNAi machinery to generate RNA guides against additional copies of the intruder RNA (Wang et al., 2006); when under a viral onslaught, maximizing cleavage rates may be more important than maximizing specificity. Moreover, the fly RNAi machinery includes additional factors and steps that may enhance specificity, including sorting siRNAs and microRNAs to different Argonaute proteins, Ago2 and Ago1 respectively (Förstemann et al., 2007; Tomari et al., 2007; Wee et al., 2012). In contrast to its fly homolog, mouse AGO2, which functions in miRNA-based targeting, exhibited dissociation and cleavage rate constants that are similar for a fully complementary target RNA (Figure 4B) (Wee et al., 2012). In this scenario, when a mismatch is introduced that increases the dissociation rate constant, the mismatched target will preferentially dissociate rather than be cleaved, providing specificity for correct targets over mismatched sequences (Figure 4B, black vs. red free energy reaction profiles). Thus, in the case of miRNA-based targeting, at least for this mouse enzyme, the enzyme kinetics may be within Regime 1, with higher intrinsic specificity than fly Ago2. Nevertheless, recent careful measurements suggest that mouse AGO2 may also be in the ‘sticky enzyme’ kinetic regime, perhaps reflecting activity at physiological temperature (Table S1) (Salomon et al., 2015). Overall, these studies of Argonaute proteins highlight the importance of extensive biochemical characterization and of interpreting and comparing results in the context of kinetic frameworks. Future comparisons of in-vitro specificity measurements with specificity determinations in vivo will be critical in discerning cellular features that impact RGN function and specificity.

### Improving RGN specificity: Lessons from biology

The previous sections highlight a paradox in RGN targeting. From a numerical perspective, uniquely targeting one sequence among many near-cognate sequences requires relatively long guides—e.g., at least ~16 nt for targeting a unique sequence in the human genome. (The probability of encountering a 16-mer of a given sequence is one in 4^16^, which is similar to the length of the human genome of ~3×10^9^ bp.) However, if a 16-nt guide bound the target with the affinity observed for Watson-Crick base pairing, the dissociation rate constant would be in the range of 10^−2^−10^−22^ s^−1^ (corresponding to GC contents ranging from 0–100%), such that dissociation would take from several minutes to 10^15^ years. For a 22 base pair duplex with the *let-7a* miRNA sequence, dissociation is predicted to take ~10^10^ years at 37 ΰC, about the lifetime of the universe! Thus, even for sequences containing multiple mismatches, with each mismatch affecting the off-rate by up to ~100-fold, there would not be sufficient time to dissociate and allow discrimination.

How has Nature addressed this challenge? Most simply, there is evidence that at least some of the guide-target base-pairing interactions are weakened in the context of an RGN. For example, experimental studies of RNAi and CRISPR targeting suggest that base pairs along the recognition sequence have differential contributions to binding and
to specificity (Ameres et al., 2007; Cong et al., 2013; Haley and Zamore, 2004; Hsu et al., 2013; Jinek et al., 2012; Jo et al., 2015; Pattanayak et al., 2013; Sternberg et al., 2014; Wee et al., 2012), and there are reports of weaker overall RISC-target interactions (Ameres et al., 2007; Haley and Zamore, 2004; Wee et al., 2012) compared to duplexes of the same sequence. An important challenge is to systematically establish quantitative rules of base-pairing in the context of RGNs, and several studies have laid the groundwork for understanding position-dependent energetic contributions (Chandradoss et al., 2015; Jo et al., 2015; Salomon et al., 2015; Sternberg et al., 2014; 2015; Wee et al., 2012). A second important challenge that follows is the determination of specificity in vivo, compared to in vitro, and identification of cellular factors that may alter specificity. For example, DNA helicases, replication, and transcription machinery may displace strongly bound CRISPR complexes and enhance specificity.

Which features of the RGN complex lead to the observed weakened binding? Most generally, a protein-bound guide is expected to be conformationally restricted, relative to a free guide strand (Bartel, 2009). In principle, such positioning can be used to enhance affinity, as is typical in macromolecular recognition, by reducing the entropic penalty for association. However, extensive protein-guide interactions may lead to conformations of the RNA guide that are suboptimal for duplex structure, and thereby reduce binding affinity relative to base-pairing in the absence of RGN.

In the case of eukaryotic Argonaute RGNs, the protein structure was noted early to be incompatible with formation of a continuous A-form duplex along the full length of the RNA guide, due to interactions with and deformations induced by the Argonaute protein (Bartel, 2009; Elkayam et al., 2012; Nakanishi et al., 2013; 2012; Schirle and MacRae, 2012; Schirle et al., 2014). Further, binding studies, analysis of miRNA-induced mRNAs repression, and the conservation of miRNA binding sites suggest that complementarity outside the short seed region of the RNA guide (positions 2-8) does not contribute strongly to recognition and repression (Doench and Sharp, 2004; Lee et al., 1993; Lewis et al., 2005; Wightman et al., 1993).

In contrast to the structures of eukaryotic Argonaute complexes, structures of Cas9 with the RNA guide and target DNA include two full helical turns (20 bp) of continuous base-pairing between the guide RNA and the target DNA, with only minor deviations from normal A-form wrapping (Anders et al., 2014; Jiang et al., 2016; Nishimasu et al., 2014). Several lines of evidence suggest that during initial probing of target sequences, this state is preceded by a conformational transition from intermediates containing only the base-pairs proximal to the protospacer adjacent motif (PAM) (Jiang et al., 2015; Sternberg et al., 2014). Conformational changes have also been suggested to accompany target recognition by Ago proteins (Nakanishi et al., 2012; Schirle et al., 2014; Sheng et al., 2014; Tomari and Zamore, 2005; Wang et al., 2009), including siRNA targeting, which, unlike miRNA targeting, relies on extensive complementarity beyond the seed region. Specificity can arise from mismatches slowing rate-limiting conformational changes. The presence of an RGN state that requires a conformational change to attain the active state will weaken overall binding and can thereby enhance specificity—i.e., the RGN reaction kinetics can be shifted from Regime 2 to Regime 1 (Fig. 3C & D). In contrast, introduction of a conformational step for an RGN already operating in Regime 1 is not expected to enhance specificity (Alber, 1981; Herschlag, 1988; Johnson, 2008).

Another strategy to improve specificity that is utilized by at least some RGNs involves increasing the rate of target *association.* Because the equilibration rate is determined by the sum of binding and dissociation rates, it follows that faster association promotes faster equilibration and, as noted above, this ability to equilibrate prior to cleavage allows intrinsic differences in binding stability to be manifest in observed specificity (Herschlag, 1991). Viewed from another perspective, the association rate constant (*k*_on_) sets the upper limit for *k*_cat_/*K*_M_ (Fersht, 1999). Thus, if cleavage was very fast, and binding was rate-limiting, specificity would be determined by the ratio of association rates for the correct versus incorrect substrate. As shown for varying association rate constants in Figure S1, faster target binding enables the system to continue to discriminate at longer guide lengths (i.e., the gray region is shifted right, to larger number of base-pairs). Consideration of association rate constants is important given that free oligonucleotides form helices orders of magnitude slower than diffusion (10^6^ M^−1^s^−1^; (Ross and Sturtevant, 1960; Wetmur and Davidson, 1968)), so that RGNs have the potential to enhance specificity by increasing the association kinetics. Indeed, Argonaute proteins have been found to accelerate target binding to the maximal diffusion controlled encounter rate (~10^8^ M^−1^s^−1^) (Jo et al., 2015; Salomon et al., 2015; Wee et al., 2012), apparently through preorganization of the seed sequence in the guide in an A-form helical conformation (Faehnle et al., 2013; Jiang et al., 2015; Nakanishi et al., 2013; Schirle and MacRae, 2012).

Finally, there are situations in which discrimination can be larger than that observed for base pair formation and binding. Especially for residues at or near the site of chemical cleavage, base-pair formation may facilitate alignment with respect to the enzyme's catalytic residues and thereby affect cleavage efficiency and contribute to specificity. This phenomenon has been described as ‘intrinsic binding energy’ by Jencks (Jencks, 1975) and has been demonstrated in ribozyme systems (Hertel et al., 1997; Narlikar et al., 1997), where base-pair contributions to cleavage rates can be greater than the thermodynamic contributions to base-pair and binding stability.

### Engineering RGNs for increased specificity

RNA-guided targeting systems evolved under different sets of selective pressures, which may have led to specificities and turnover rates optimized for a particular natural context. Thus, it may not be surprising that initial efforts to apply these systems for laboratory applications revealed specificities that fall short of those required to, for example, target unique sites in mammalian transcriptomes or genomes (Hu et al., 2016; Jackson et al., 2003; Wu et al., 2014). One straightforward reason for this apparent shortcoming was introduced above: natural systems, such as fly Ago2, and, likely, the bacterial CRISPR machinery may simply not have evolved to be exquisitely specific, instead responding to the evolutionary pressure to mount a rapid and efficacious response against foreign nucleic acids. Minimizing off-target effects may not be a major selective pressure for bacterial RGNs (such as CRISPR-Cas9), in part because of the small sizes of bacterial genomes: the probability of encountering detrimental off-targets is lower in a genome that is only ~1 Mb, the size of the *Streptococcus pyogenes* genome, which is the source of the widely used Cas9 protein, than it is in the 3 Gb human genome. Developing RGN systems to be highly specific tools may therefore require modifying their inherent affinities and reaction rates by engineering or selection.

In this section, we present two general strategies for improving specificity based on the kinetic considerations laid out in the previous sections. The unifying premise of these strategies is that the ability of an RGN to discriminate between correct and mismatched targets increases as dissociation becomes more favorable than cleavage. As was described above, discrimination can be considered in the free energy diagram in Figure 3 in terms of the relative heights of the two peaks for dissociation and cleavage that lead from the E•S complex (Figure 3). Thus, it follows that discrimination will increase upon increasing *k*_off_ or decreasing *k*_cat_, and the greatest discrimination will be achieved upon simultaneously changing both rate constants. Indeed, many of the successful strategies reported in the literature appear to follow these strategies (Table 1). In a few of these cases, on-target activity, such as gene repression, appears to be maintained when destabilizing modifications are made, while off-targeting decreases (Hu et al., 2016; Jackson et al., 2006; Ohnishi et al., 2008; Pfister et al., 2009). These effects would not occur if thermodynamics alone were at play (Figure 2B), implying the importance of kinetic considerations for the specificity of these systems (Figure 2C).

**Table 1.**
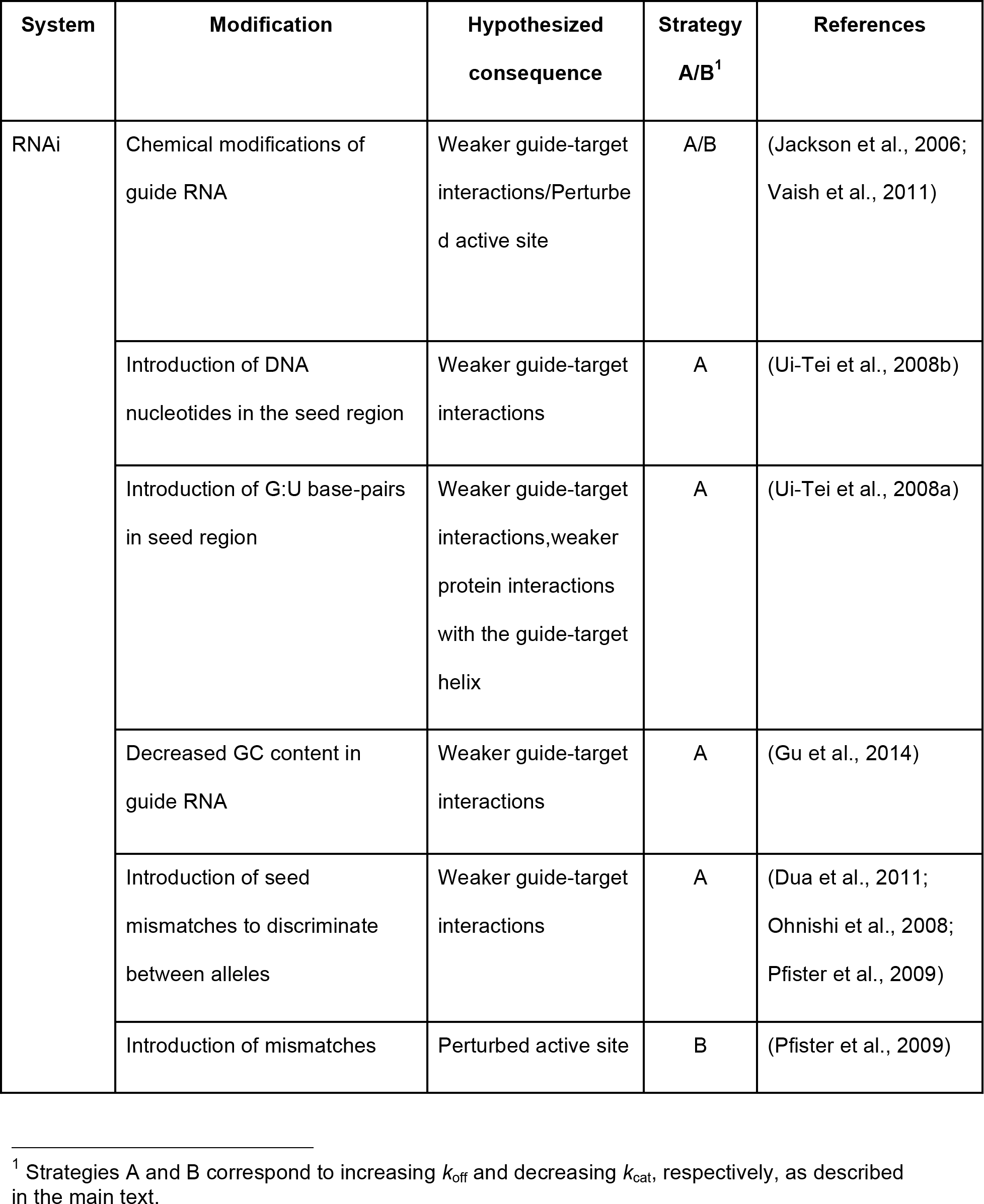
Strategies to improve specificity of RNA-guided targeting

|  |  |  |  |  |
| --- | --- | --- | --- | --- |
|  | near the catalytic site |  |  |  |
| CRISPR-Cas9 | Truncated guide RNA | Weaker guide-target interactions | A | (Fu et al., 2014; Tsai et al., 2015) |
|  | Cas9 mutations neutralizing positive charge in the DNA binding grooves | Weaker Cas9-target interactions | A | (Kleinstiver et al., 2016; Slaymaker et al., 2016) |
| RNaseH<sup>2</sup> | Chemical modifications of guide DNA | Perturbed active site | B | (Østergaard et al., 2013) |

<sup>2</sup> RNase H is technically a DNA-guided nuclease that cleaves the complementary RNA target. We include it in our discussions because it is conceptually similar to and has been utilized in similar fashion to the RGNs discussed herein.

#### Strategy A: Increasing *k*_off_

The RGN literature is rich in examples of increased specificity upon perturbing RGN-target interactions, and many of these examples can be explained by increased *k*_off_ (Table 1). The most straightforward strategy to weaken target-guide interactions involves shortening the guide to decrease the number of base-pairs formed with the target sequence.

For example, truncated CRISPR guide RNAs showed reduced off-target activity, in some cases—with no observed reduction of cleavage of the intended target (Fu et al., 2014; Tsai et al., 2015). This observation is consistent with a switch from the “sticky” to Michaelis-Menten regime (Figure 5, Strategy A), resulting in enhanced specificity. Although in practice RNA guides can only be shortened by a few bases—both to maintain sufficient number of contacts for robust on-target activity and because recognizing individual genomic sites imposes minimum length requirements on the guide—even truncating guide RNAs by 2-3 nucleotides has provided substantial specificity enhancements in CRISPR-Cas9 targeting (Fu et al., 2014). These and other results suggest that destabilizing binding actually increases specificity of S. *pyogenes* Cas9 and that this system functions as a “sticky” enzyme in human cells (Figure 3D).

**Figure 5.**
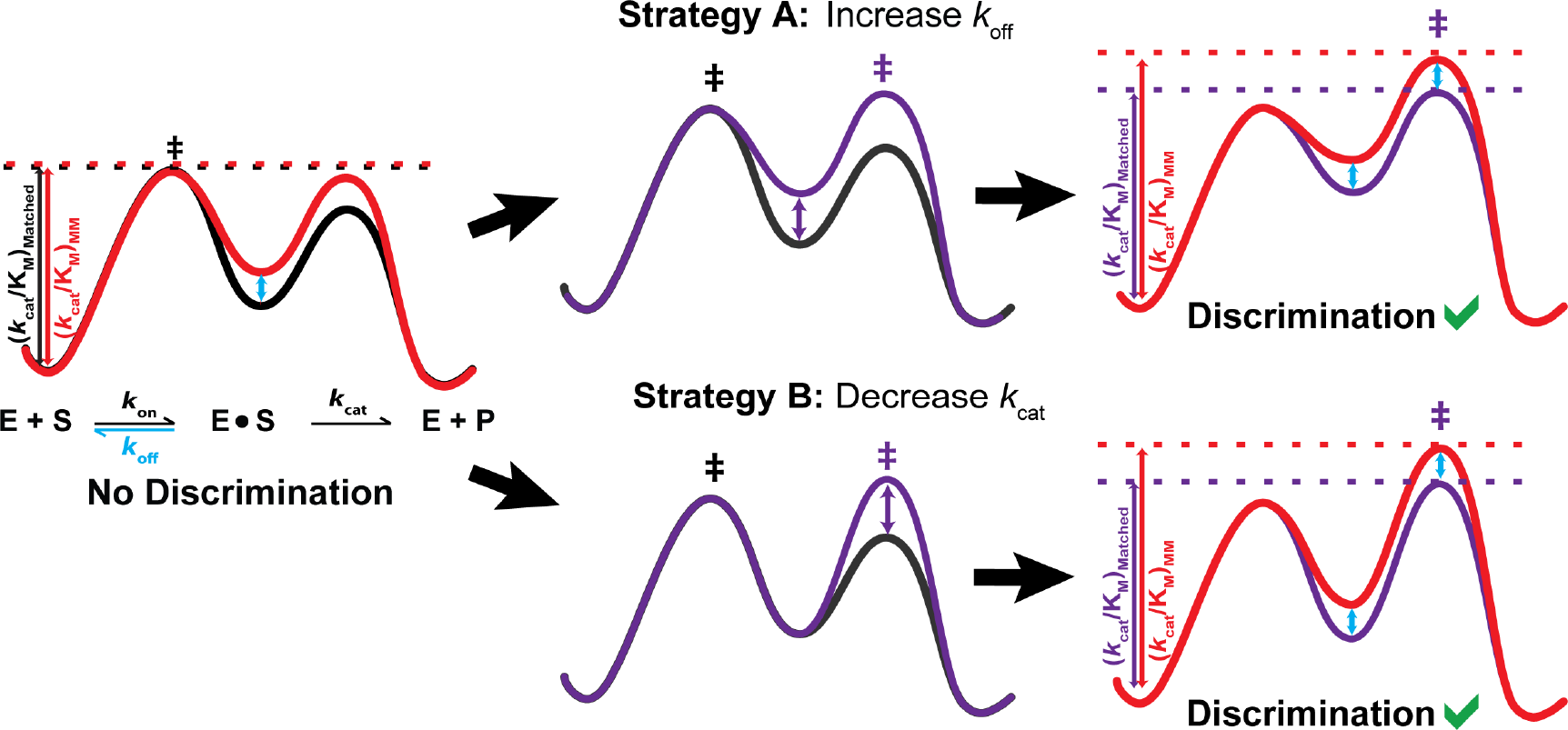
Strategies to increase specificity. **Left.** Free energy profile representing the kinetics of a“sticky” enzyme for a fully complementary (black) and a mismatched target (red). The efficiency of cleavage, *k*_c_/*K*_m_, is designated with vertical lines to the transition state (‡) of the reaction. Because dissociation is slow relative to cleavage, the efficiency of both reactions is the same and there is no specificity. The free energy diagram depicts a scenario in which the association rate (*k*_on_) and cleavage rate (*k*_cat_ are unchanged for the mismatched target, while the dissociation rate (*k*_off_) is faster (blue arrow). **Middle**. Two strategies to increase discrimination, change the rate-limiting step from binding to cleavage (changes shown for the complementary target only). Reaction profiles are shown before (black) and after a change in the rate-limiting step (purple). **Right**. New free energy diagrams for the modified reactions of the matched (purple) and mismatched (red) target with the off-rate increased (top) or the cleavage rate decreased (bottom). Modifications result in differences in *k*_cat_/*K*_M_ (blue arrow) between the two substrates and thus enable discrimination.

Target binding can be weakened without sacrificing the number of residues recognized, by lowering the average base-pair stability. As was originally described in the context of ribozyme targeting (Herschlag, 1991), the decreased average stability of a base-pair (e.g., in an AU-rich vs. a GC-rich sequence) leads to a higher threshold for discrimination in a random pool of target sequences, as more residues can be used in recognition before dissociation becomes slower than cleavage of mismatched targets. This concept can be extended to targeting by any RGN: for instance, specificity improvements in RNAi were observed after increasing the AU content in the guide (Gu et al., 2014). We note that the effects of increased AU content may be especially large in the case of DNA-targeting RGNs that use RNA guides because of the exceptionally low stability of dA-rU base-pairs (Martin and Tinoco, 1980). In certain cases, the weaker stability of DNA/RNA compared to RNA/RNA base pairs (Ui-Tei et al., 2008b) and the additional destabilization of dA/rU pairs might be used to enhance specificity. Certain chemical modifications of the guide could also destabilize target binding, by lowering base-pair stability and/or weakening contacts between target and the protein.

Finally, target binding can be weakened by introducing mutations into the nuclease itself. Significant improvements in CRISPR-Cas9 specificity have been achieved by substitutions of positively charged residues in the ssDNA binding sites of Cas9, hypothesized to weaken electrostatic interactions with the DNA backbone (Kleinstiver et al., 2016; Slaymaker et al., 2016).

#### Strategy B: Decreasing *k*_cgt_

Decreasing the rate of target cleavage can increase the specificity of a “sticky” enzyme by promoting binding equilibration prior to cleavage (Figure 5, Strategy B). Although a decrease in on-target cleavage is generally viewed as a flaw and corresponding mutations and constructs are dismissed, it may be fruitful to reexamine nuclease and/or guide variants that give slower cleavage to optimize specificity.

How can *k*_cat_ be manipulated? In vitro measurements have shown that mismatches in the miRNA-target duplex in proximity of the cleavage site reduce *k*_cat_ (Wee et al., 2012), and the same likely applies to other RGNs, presumably as a result of active site perturbations. Chemical modifications of the guide have been reported to improve RNAi specificity, and some of these modifications may primarily affect *k*_cat_ rather than the dissociation rate, or may affect both (Jackson et al., 2006; Vaish et al., 2011; Østergaard et al., 2013). Finally, nuclease domains with varying cleavage rates may be tethered to a catalytically inactive RGN to achieve a range of activities.

There will be a point at which these manipulations will make cleavage too slow to act on the required timescale, or dissociation will be too fast to efficiently target the desired molecules. Thus, there will be a kinetic 'sweet spot’ for a given RGN and application that can be identified through systematic studies. Moreover, some manipulations could potentially introduce detrimental effects on other steps (such as Cas9 or Argonaute loading with the guide RNA), which will need to be accounted for. Given these complexities, conceptual strategies such as those presented here are needed to guide rational re-engineering of RGNs, to identify the limits of improved RGN specificity, and to determine the *in vivo* factors that adjust RGN reaction properties.

#### Other strategies

Other successful strategies for improving RGN specificity have been reported and we refer the reader to a recent review of these strategies (Hu et al., 2016). Here we briefly note several of these approaches to place them in the context of basic kinetic and thermodynamic principles, analogous to the discussions above.

Improvements in specificity have been observed with lower RGN concentrations: i.e., less production of the RGN complex or when RGN activity was controlled with small molecule activators and repressors (reviewed in (Hu et al., 2016)). This may arise from very high RGN concentrations that exceed the amount of substrate and the reaction's *K*_M_ values, such that targets and off-targets may no longer need to compete for a common limiting pool of RGNs complexes (Herschlag, 1988).

Another approach that has been noted to improve specificity is limiting the time of the RGN reaction (Hu et al., 2016). In much of RGN literature ‘observed specificity’ corresponds to the relative *amounts* of target versus off-target cleaved at a given sampling time point, rather than relative *rates* of the reaction. But the cleavage of all possible targets will go to completion at some point if a reaction is allowed to proceed for a sufficient time, with slower-cleaving off-targets ‘catching up’ to targets and the apparent discrimination lessening over time (Figure S2). Thus, it is essential to monitor time courses of cleavage, rather than sampling at an arbitrary single time point, and to adjust the time of cleavage and the concentration of the RGN accordingly.

### In practice: Strategies for discrimination between single-nucleotide polymorphisms

RGNs have long been recognized as promising therapeutics because of their potential to specifically target mutated genes associated with disease, especially autosomal dominant disorders. Preclinical data exist for RNAi targeting of disease-associated single-nucleotide polymorphisms (SNPs) of hereditary diseases, such as Huntington's disease, hypertrophic cardiomyopathy, amyotrophic lateral sclerosis, and sickle-cell anemia (Dykxhoorn et al., 2006; Jiang et al., 2013; Miller et al., 2003; Pfister et al., 2009; Schwarz et al., 2006; Yu et al., 2012; Østergaard et al., 2013), and there are promising studies of gene editing using CRISPR-Cas9 systems of human hereditary diseases (Porteus, 2016; Yin et al., 2016). In these RNAi studies, the therapeutic siRNA is designed to have full complementarity to the disease allele and to contain a mismatch to the wildtype allele at the site of disease-associated polymorphism (Figure 6A). A key challenge is that, since disease alleles often differ from the wild-type allele by only a single nucleotide, there is often little or no discrimination in silencing between wild-type and disease alleles.

**Figure 6.**
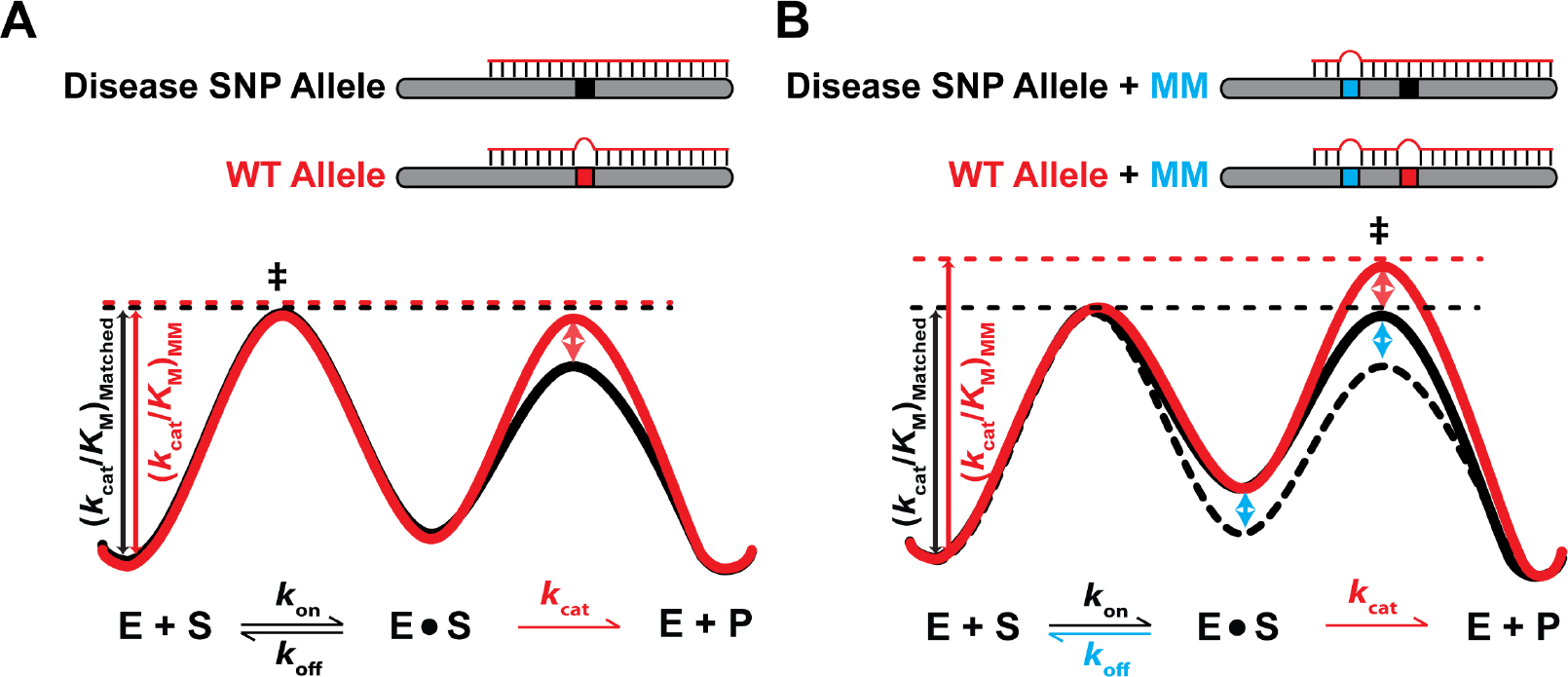
Discrimination strategies for single-nucleotide polymorphisms and energetic ‘threshold’ effects. **A**. An siRNA that is fully complementary to the disease allele (black) contains a mismatch against the WT allele at the location of the SNP (red). This mismatch is often positioned in the central region of the siRNA (nucleotides 9-11) to maximize specificity. The effect of a central mismatch is expected to be on the cleavage rate, *k*_cat_(Nee et al., 2012), as reflected in the free energy diagram for the WT allele (red) and the disease allele (black). Because the rate-limiting step is still binding for both the WT and the disease alleles, there is no (or little) discrimination in silencing (Regime 2, Figure 3D). **B**. Introduction of an additional seed mismatch (blue) for both the disease and WT allele. The additional mismatch destabilizes binding (blue arrow) of both the WT (red) and disease allele (black) targets relative to binding with only a single mismatch (dotted black line), and binding to the WT allele is further destabilized by the mismatch at the SNP location. This additional destabilization leads to a change in *k*_cat_/*K*_M_, and thus an increase in discrimination between the two alleles. This additional destabilization allows the system to cross the threshold from Regime 2 to Regime 1 (Figure 3C&D).

Researchers have determined empirically that the largest discrimination in the efficiency of siRNA-mediated silencing often occurs when the mismatch against the WT allele is located in the central region of the guide (positions 9-11), which lies near the cleavage site of Argonaute and can have large effects on the cleavage reaction (Wee et al., 2012). Introduction of an additional mismatch in the seed region of the guide against *both* the WT and disease allele has also been shown to increase specificity—i.e., these constructs have greater discrimination than for each mismatch alone (Dahlgren et al., 2008; Jiang et al., 2013; Miller et al., 2003; Pfister et al., 2009). These observations can be explained by a threshold effect due to a combination of strategies A and B (Narlikar and Herschlag, 1998; Narlikar et al., 1997): the mismatch at the central residues reduces *k*_cat_ (Strategy B (Ameres et al., 2007; Haley and Zamore, 2004; Wee et al., 2012)) and the mismatch in the seed region increases *k*_off_ (Strategy A; (Ameres et al., 2007; Haley and Zamore, 2004; Wee et al., 2012)), but neither effect is sufficient to allow discrimination (Regime 2 is maintained; Figure 6A). When both changes are made together, a 'threshold’ is crossed from Regime 2 to Regime 1, as illustrated in Figure 6B relative to Figure 6A.

### Summary and perspective

RGN systems hold tremendous technological and therapeutic promise, and recent advances in increasing their delivery, stability, and specificity have markedly lowered the barriers to wide application of these potent and versatile tools (Haussecker and Kay, 2015; Hendel et al., 2015; Hu et al., 2016; Wittrup and Lieberman, 2015). Here we approached the widespread problem of off-target effects in RGN targeting from a kinetic perspective that explains and codifies previous observations of RGN specificity in terms of simple and generalizable models and offers rational strategies for minimizing off-target activity.

As we highlight, the most obvious ‘culprit’ in the widespread inability of RGN systems to efficiently discriminate between targets and very similar sequences is the inherent high stability of base-pairing interactions that target recognition relies on. Very slow dissociation of stable target-guide duplexes can lead to a kinetic regime where the RGN cleaves fully matched and mismatched targets before having an opportunity to discriminate between them based on differences in dissociation rates. This limitation is alleviated, at least in part, by RGN proteins that destabilize guide-target interactions by deforming the duplex and blocking helix propagation along the full length of the guide. An effective general strategy to improve RGN specificity is through further weakening of the RGN-target interactions, and here we have reevaluated published strategies that take this approach in terms of the underlying kinetic steps and presented additional rational approaches guided by kinetic models of specificity. Some of the strategies unearthed in studies of RNAi may also apply to CRISPR and vice versa.

While we chose to use a minimal two-step model of RGN activity for the purpose of explaining kinetic concepts, we emphasize that the ideas and kinetic regimes presented here naturally extend to more complex scenarios. For example, *k*_cat_ can represent any downstream rate-limiting transition of interest in a complex reaction scheme. This may include steps such as conformational changes that have been identified in several cases (Jiang et al., 2015; Sternberg et al., 2015), product release (Haley and Zamore, 2004; Salomon et al., 2015), and downstream steps such as mRNA turnover that likely follows miRNA-dependent translation inhibition (Larsson et al., 2010). Moreover, additional in vivo factors could facilitate increases in off-rates, such as helicases and polymerases that might dislodge CRISPR complexes, and ribosomes that might dislodge RISC complexes. More generally, the natural environments in which RGN systems evolved to operate may contribute to specificity, and understanding these contributions will be important for maximizing the potency and specificity of RGNs in new contexts.

While it might appear at first glance that the complexities of RGN systems and the cellular milieu render attempts to describe basic principles hopelessly naïve, we believe just the opposite to be true. To understand and ultimately tame a complex system, a stepwise approach is required. For RGNs, identifying their reaction steps and understanding their thermodynamics and kinetics in isolation will allow firm and quantitative predictions to be made and tested in vivo, potentially leading to the identification of new in vivo factors that can ultimately be exploited for engineering and medicine.

To our knowledge, cleavage and dissociation kinetics are currently not taken into account in any prediction algorithm for RGN target-sites and off-target effects. The energetics of base-pairing interactions in the context of RGNs remains to be fully established, and also how off-target dissociation rates compare to cleavage rates in natural and engineered RGN systems. The extensive, quantitative studies of RGN systems (Chandradoss et al., 2015; De et al., 2013; Haley and Zamore, 2004; Jo et al., 2015; Salomon et al., 2015; Sternberg et al., 2014; Wee et al., 2012) inaugurate the types of analyses that will allow physical models to be developed and predictions to be made for specificity and efficacy. We hope that this perspective will stimulate further quantitative, mechanistic studies of RGNs, which in turn will enable more accurate predictions and rational approaches to RGN re-engineering and application.

## MATERIALS AND METHODS

<u>Simulation of binding and cleavage reactions.</u> For Figure 1, thermodynamic stabilities of *let-7a* guide RNA to RNA targets of varying length were calculated using the Dinamelt Web Server (Markham and Zuker, 2005). The default solution conditions were used at 37 °C. Binding affinities were calculated from these ΔG predictions using ΔG = - RTln(*K*_A_). Kinetic simulations were performed using *k*_on_ = 4×10^7^M^−1^s^−1^ for the binding of the RGN-guide RNA complex to the target (based on a measurement for mouse AGO2 from (Wee et al., 2012)). Off-rates were inferred from *K*_A_ = *k*_on_/*k*_off_. Specificity in Figure 1C was determined from the ratio of the *k*_cat_/*K*_M_ values for the fully complementary target vs. each of the mismatched targets. For illustration purposes, *k*_cat_ was assumed to be 10′ ^8^s^−1^, which is likely to be slower than physiological rates, but faster rates will only exacerbate decreases in specificity with increasing guide length. Simulations with different values of these parameters are shown in Figure S1.

<u>Determination of dissociation times for 16 bp RNA:RNA duplexes:</u> The range of dissociation times was calculated based on the stabilities of an AU-only versus a GC-only 16 bp RNA:RNA helix. The half-life, or time required for 50% of helices to dissociate, equals: t_1/2_ = ln(2)/*k*_off_, and *k*_off_ is derived from the stability modeled by Dinamelt (Markham and Zuker, 2005) at 37 °C: *k*_off_= *k*_on_/*K*_eq_ = *k*_on_/e^−ΔG/RT^, assuming *k*_on_ = 10^6^M^−1^s^−1^ for duplex formation (Ross and Sturtevant, 1960; Wetmur and Davidson, 1968). Dissociation times are reported as five half-lives.

## ACKNOWLEDGEMENTS

We thank Liang Wee, members of the Herschlag and Bartel labs, Matt Porteus, Andy Fire, Stanley Qi and Mark Kay for helpful discussions, comments, and for critical advice throughout this work. N.B. was supported by an NSF graduate fellowship, and this work was supported by a grant to D.H. from the National Institutes of Health [P01GM066275].

## SUPPLEMENTAL INFORMATION

**Figure S1.**
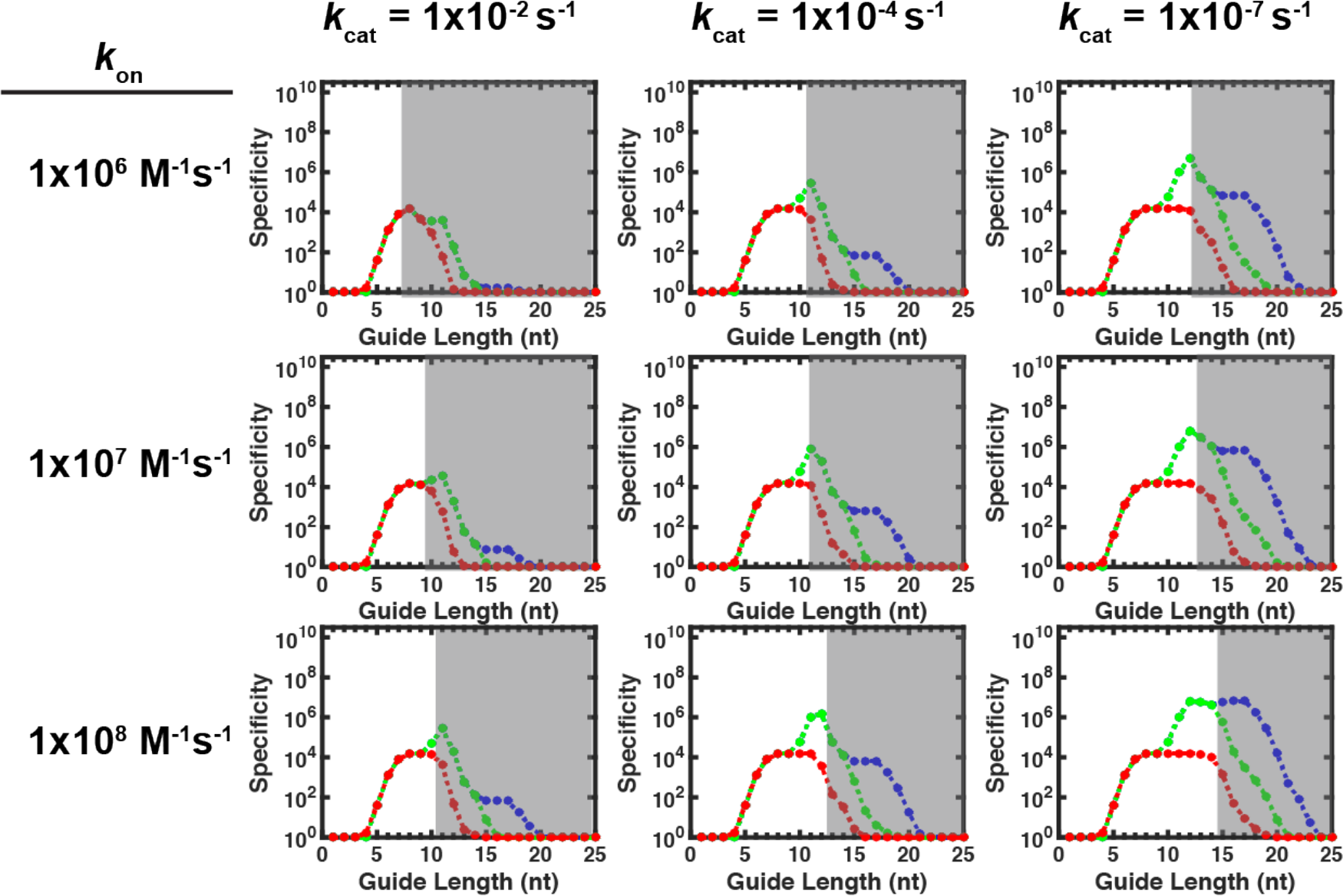
Effect of varying *k*_on_ and *k*_cat_ values on specificity. The simulations are analogous to that in Figure 2C, except for using different *k*_on_ and /*f*_cat_ values. Specificity is defined as the ratio of *kcJKu* values for the fully complementary target relative to off-targets containing one, two, or three mismatches (red, green, and blue lines, respectively). Each row uses a different specified on-rate (*k*_on_), with values shown on the left, and each column uses a different cleavage rate constant (*k*_cat_), denoted on top. Dissociation rate constants in the simulation were the same in each panel, and taken from Figure 2 (see Methods). Gray areas denote regions where thermodynamic specificity diverges from kinetic specificity.

**Figure S2.**
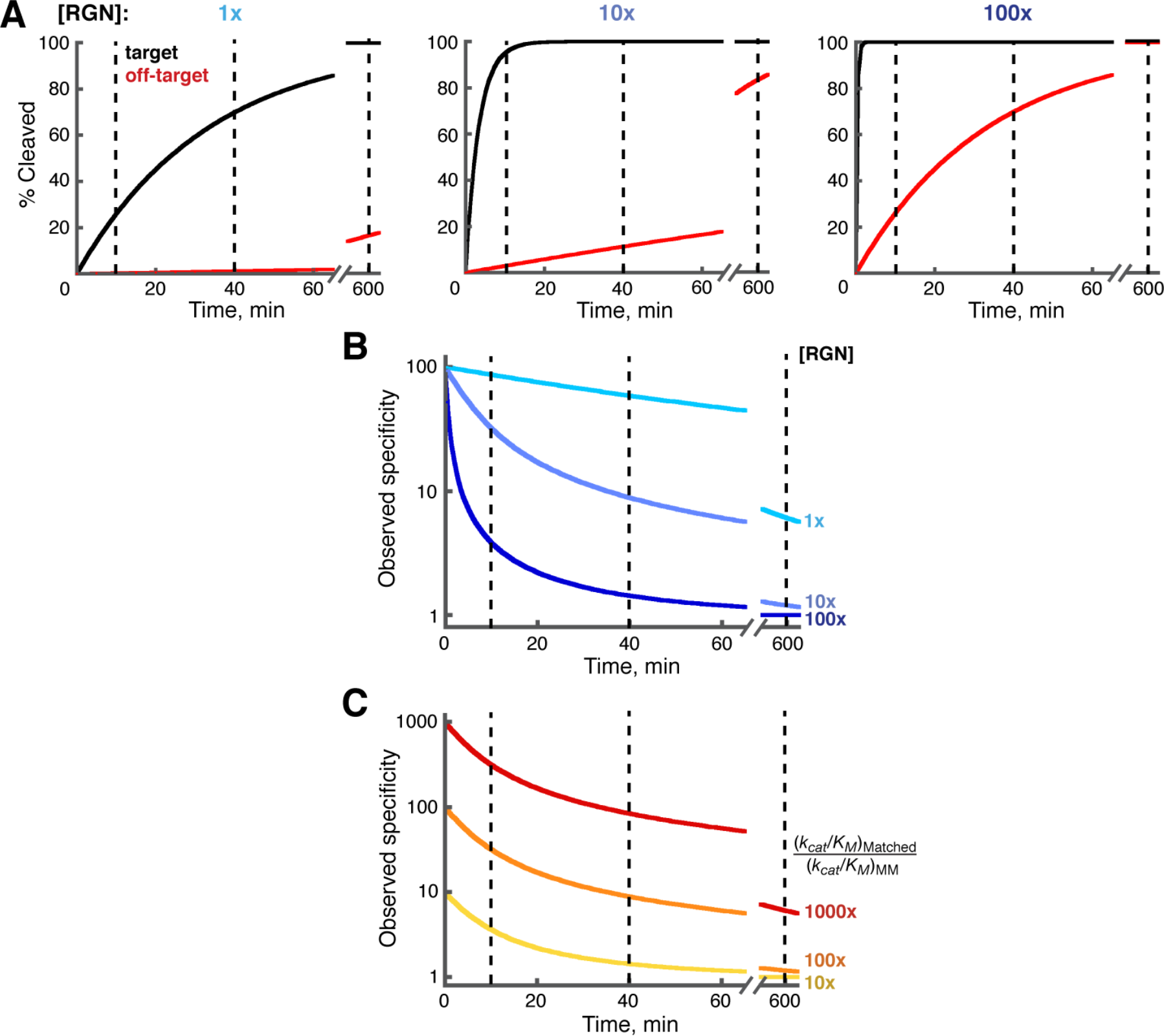
Duration of the RGN reaction affects ‘observed specificity’. **A**. Cleavage time courses of a hypothetical RGN at three RGN concentrations (1x, 10x, 100x). The target (black) is always cleaved at a 100-fold greater rate than the off-target (red), corresponding to the ratio of *k*_ca_/*K*_M_ values for target vs. off-target cleavage. Dashed lines mark three representative time points along the reaction time course. Note that the difference between the amount of target vs. off-target cleaved decreases as the target is depleted over time, until both target and off-target are essentially completely cleaved at long times and high RGN concentrations. **B**. Changes in ‘observed specificity’ over time. Observed specificity is defined as the ratio of cleaved fractions of target vs. off-target at a given time. The colors correspond to the RGN concentrations in (A). Note that the observed specificity decreases over time, and it decreases more rapidly at higher RGN concentrations. **C**. Time-dependent decline in observed RGN specificity for RGN systems with different inherent specificities, as defined by the ratio of *k*_cat_/*K*_M_ values for target vs. off-target. The colors correspond to *k*_ca_/K_M_ ratios of 1000-(red), 100-(orange) and 10-fold (yellow). Note that observed discrimination becomes more similar in all cases at longer times; sampling the reaction at a single time point (e.g., 10 hours) can result in lack of observed specificity, regardless of the extent of inherent specificity.

**Table S1.** Rate constants for fly Ago2 and mouse AGO2 in complex with let-7a guide RNA and fully complementary target RNA

| Enzyme | Temperature, °C | Reference | $k_{off}$ (s<sup>-1</sup>)<sup>a</sup> | $k_{cat}$ (s<sup>-1</sup>) |
| --- | --- | --- | --- | --- |
| Fly Ago2 | 25 | (Wee et al., 2012) | $8.8 \times 10^{-5}$ | $6.1 \times 10^{-2}$ |
| Mouse AGO2 | 25 | (Wee et al., 2012) | $7.7 \times 10^{-4}$ | $8.1 \times 10^{-4}$ |
| | 37 | (Salomon et al., 2015) | $\leq 3.6 \times 10^{-3}$<sup>b</sup> | $7.8 \times 10^{-2}$ |

<sup>a</sup> Dissociation measurements at 25 °C were carried out with targets containing 2'-O-methyl ribose flanked by phosphorothioate linkage at the scissile site to prevent cleavage.
<sup>b</sup> The dissociation rate constant was not measured for the fully complementary target at 37 °C; the dissociation from a target in which complementarity is limited to the seed region provides an upper estimate of the dissociation rate constant (Salomon et al., 2015).

## REFERENCES

Alber, T.C. (1981). Structural origins of the catalytic power of triose phosphate isomerase. PhD Thesis, Massachusetts Institute of Technology.

Ameres, S.L., Martinez, J., and Schroeder, R. (2007). Molecular basis for target RNA recognition and cleavage by human RISC. Cell 130, 101–112.

Anders, C., Niewoehner, O., Duerst, A., and Jinek, M. (2014). Structural basis of PAM-dependent target DNA recognition by the Cas9 endonuclease. Nature 513, 569–573.

Bartel, D.P. (2009). MicroRNAs: target recognition and regulatory functions. Cell 136, 215–233.

Briggs, G.E., and Haldane, J.B. (1925). A note on the kinetics of enzyme action. Biochem. J. 19, 338–339.

Carthew, R.W., and Sontheimer, E.J. (2009). Origins and mechanisms of miRNAs and siRNAs. Cell 136, 642–655.

Chandradoss, S.D., Schirle, N.T., Szczepaniak, M., MacRae, I.J., and Joo, C. (2015). A dynamic search process underlies MicroRNA targeting. Cell 162, 96–107.

Cong, L., Ran, F.A., Cox, D., Lin, S., Barretto, R., Habib, N., Hsu, P.D., Wu, X., Jiang, W., Marraffini, L.A., et al. (2013). Multiplex genome engineering using CRISPR/Cas systems. Science 339, 819–823.

Dahlgren, C., Zhang, H.-Y., Du, Q., Grahn, M., Norstedt, G., Wahlestedt, C., and Liang, Z. (2008). Analysis of siRNA specificity on targets with double-nucleotide mismatches. Nucleic Acids Res *36*, e53.

De, N., Young, L., Lau, P.-W., Meisner, N.-C., Morrissey, D.V., and MacRae, I.J. (2013). Highly complementary target RNAs promote release of guide RNAs from human Argonaute2. Mol Cell 50, 344–355.

Doench, J.G., and Sharp, P.A. (2004). Specificity of microRNA target selection in translational repression. Genes Dev 18, 504–511.

Doench, J.G., Fusi, N., Sullender, M., Hegde, M., Vaimberg, E.W., Donovan, K.F., Smith, I., Tothova, Z., Wilen, C., Orchard, R., et al. (2016). Optimized sgRNA design to maximize activity and minimize off-target effects of CRISPR-Cas9. Nat Biotechnol 34, 184–191.

Dominguez, A.A., Lim, W.A., and Qi, L.S. (2016). Beyond editing: repurposing CRISPR-Cas9 for precision genome regulation and interrogation. Nat Rev Mol Cell Bio 17, 5–15.

Dua, P., Yoo, J.W., Kim, S., and Lee, D.-K. (2011). Modified siRNA structure with a single nucleotide bulge overcomes conventional siRNA-mediated off-target silencing. Mol. Ther. 19, 1676–1687.

Dykxhoorn, D.M., Schlehuber, L.D., London, I.M., and Lieberman, J. (2006). Determinants of specific RNA interference-mediated silencing of human beta-globin alleles differing by a single nucleotide polymorphism. Proc Natl Acad Sci USA 103, 5953–5958.

Elkayam, E., Kuhn, C.-D., Tocilj, A., Haase, A.D., Greene, E.M., Hannon, G.J., and Joshua-Tor, L. (2012). The structure of human argonaute-2 in complex with miR-20a. Cell 150, 100–110.

Faehnle, C.R., Elkayam, E., Haase, A.D., Hannon, G.J., and Joshua-Tor, L. (2013). The making of a slicer: activation of human Argonaute-1. Cell Rep 3, 1901–1909.

Fersht, A. (1999). Structure and mechanism in protein science : a guide to enzyme catalysis and protein folding (New York: W.H. Freeman).

Förstemann, K., Horwich, M.D., Wee, L., Tomari, Y., and Zamore, P.D. (2007). Drosophila microRNAs are sorted into functionally distinct argonaute complexes after production by dicer-1. Cell 130, 287–297.

Fu, Y., Sander, J.D., Reyon, D., Cascio, V.M., and Joung, J.K. (2014). Improving CRISPR-Cas nuclease specificity using truncated guide RNAs. Nat Biotechnol 32, 279–284.

Gu, S., Zhang, Y., Jin, L., Huang, Y., Zhang, F., Bassik, M.C., Kampmann, M., and Kay, M.A. (2014). Weak base pairing in both seed and 3’ regions reduces RNAi off-targets and enhances si/shRNA designs. Nucleic Acids Res 42, 12169–12176.

Guilinger, J.P., Pattanayak, V., Reyon, D., Tsai, S.Q., Sander, J.D., Joung, J.K., and Liu, D. R. (2014). Broad specificity profiling of TALENs results in engineered nucleases with improved DNA-cleavage specificity. Nat Methods 11, 429–435.

Haley, B., and Zamore, P.D. (2004). Kinetic analysis of the RNAi enzyme complex. Nat Struct Mol Biol 11, 599–606.

Haussecker, D., and Kay, M.A. (2015). RNA interference. Drugging RNAi. Science 347, 1069–1070.

Hendel, A., Bak, R.O., Clark, J.T., Kennedy, A.B., Ryan, D.E., Roy, S., Steinfeld, I., Lunstad, B.D., Kaiser, R.J., Wilkens, A.B., et al. (2015). Chemically modified guide RNAs enhance CRISPR-Cas genome editing in human primary cells. Nat Biotechnol 33, 985–989.

Herschlag, D. (1988). The role of induced fit and conformational changes of enzymes in specificity and catalysis. Bioorg Chem 16, 62–96.

Herschlag, D. (1991). Implications of ribozyme kinetics for targeting the cleavage of specific RNA molecules in vivo: more isn't always better. Proc Natl Acad Sci USA 88, 6921–6925.

Hertel, K.J., Peracchi, A., Uhlenbeck, O.C., and Herschlag, D. (1997). Use of intrinsic binding energy for catalysis by an RNA enzyme. Proc Natl Acad Sci USA 94, 84978502.

Hsu, P.D., Scott, D.A., Weinstein, J.A., Ran, F.A., Konermann, S., Agarwala, V., Li, Y., Fine, E.J., Wu, X., Shalem, O., et al. (2013). DNA targeting specificity of RNA-guided Cas9 nucleases. Nat Biotechnol 31, 827–832.

Hu, J.H., Davis, K.M., and Liu, D.R. (2016). Chemical Biology Approaches to Genome Editing: Understanding, Controlling, and Delivering Programmable Nucleases. Cell Chemical Biology 23, 57–73.

Jackson, A.L., Bartz, S.R., Schelter, J., Kobayashi, S.V., Burchard, J., Mao, M., Li, B., Cavet, G., and Linsley, P.S. (2003). Expression profiling reveals off-target gene regulation by RNAi. Nat Biotechnol 21, 635–637.

Jackson, A.L., Burchard, J., Leake, D., Reynolds, A., Schelter, J., Guo, J., Johnson, J.M., Lim, L., Karpilow, J., Nichols, K., et al. (2006). Position-specific chemical modification of siRNAs reduces “off-target” transcript silencing. RNA 12, 1197–1205.

Jencks, W.P. (1975). Binding energy, specificity, and enzymatic catalysis: the circe effect. Adv Enzymol Ramb 43, 219–410.

Jiang, F., Taylor, D.W., Chen, J.S., Kornfeld, J.E., Zhou, K., Thompson, A.J., Nogales, E., and Doudna, J.A. (2016). Structures of a CRISPR-Cas9 R-loop complex primed for DNA cleavage. Science 351, 867–871.

Jiang, F., Zhou, K., Ma, L., Gressel, S., and Doudna, J.A. (2015). A Cas9-guide RNA complex preorganized for target DNA recognition. Science 348, 1477–1481.

Jiang, J., Wakimoto, H., Seidman, J.G., and Seidman, C.E. (2013). Allele-specific silencing of mutant Myh6 transcripts in mice suppresses hypertrophic cardiomyopathy. Science 342, 111–114.

Jinek, M., Chylinski, K., Fonfara, I., Hauer, M., Doudna, J.A., and Charpentier, E. (2012). A programmable dual-RNA-guided DNA endonuclease in adaptive bacterial immunity. Science 337, 816–821.

Jo, M.H., Shin, S., Jung, S.-R., Kim, E., Song, J.-J., and Hohng, S. (2015). Human Argonaute 2 has diverse reaction pathways on target RNAs. Mol Cell 59, 117–124.

Johnson, K.A. (2008). Role of induced fit in enzyme specificity: a molecular forward/reverse switch. J Biol Chem 283, 26297–26301.

Kleinstiver, B.P., Pattanayak, V., Prew, M.S., Tsai, S.Q., Nguyen, N.T., Zheng, Z., and Joung, J.K. (2016). High-fidelity CRISPR-Cas9 nucleases with no detectable genome-wide off-target effects. Nature 529, 490–495.

Larsson, E., Sander, C., and Marks, D. (2010). mRNA turnover rate limits siRNA and microRNA efficacy. Mol Syst Biol 6, 433.

Lee, R.C., Feinbaum, R.L., and Ambros, V. (1993). The C. elegans heterochronic gene lin-4 encodes small RNAs with antisense complementarity to lin-14. Cell 75, 843–854.

Lewis, B.P., Burge, C.B., and Bartel, D.P. (2005). Conserved seed pairing, often flanked by adenosines, indicates that thousands of human genes are microRNA targets. Cell 120, 15–20.

Mali, P., Aach, J., Stranges, P.B., Esvelt, K.M., Moosburner, M., Kosuri, S., Yang, L., and Church, G.M. (2013). CAS9 transcriptional activators for target specificity screening and paired nickases for cooperative genome engineering. Nat Biotechnol 31, 833–838.

Markham, N.R., and Zuker, M. (2005). DINAMelt web server for nucleic acid melting prediction. Nucleic Acids Res 33, W577–W581.

Martin, F.H., and Tinoco, I. (1980). DNA-RNA hybrid duplexes containing oligo(dA:rU) sequences are exceptionally unstable and may facilitate termination of transcription. Nucleic Acids Res 8, 2295–2299.

Michaelis, L., Menten, M.L., Johnson, K.A., and Goody, R.S. (2011). The original Michaelis constant: translation of the 1913 Michaelis-Menten paper. Biochemistry 50, 8264–8269.

Miller, V.M., Xia, H., Marrs, G.L., Gouvion, C.M., Lee, G., Davidson, B.L., and Paulson, H.L. (2003). Allele-specific silencing of dominant disease genes. Proc Natl Acad Sci USA 100, 7195–7200.

Nakanishi, K., Ascano, M., Gogakos, T., Ishibe-Murakami, S., Serganov, A.A., Briskin, D., Morozov, P., Tuschl, T., and Patel, D.J. (2013). Eukaryote-specific insertion elements control human ARGONAUTE slicer activity. Cell Rep 3, 1893–1900.

Nakanishi, K., Weinberg, D.E., Bartel, D.P., and Patel, D.J. (2012). Structure of yeast Argonaute with guide RNA. Nature 486, 368–374.

Narlikar, G.J., and Herschlag, D. (1998). Direct demonstration of the catalytic role of binding interactions in an enzymatic reaction. Biochemistry 37, 9902–9911.

Narlikar, G.J., Khosla, M., Usman, N., and Herschlag, D. (1997). Quantitating tertiary binding energies of 2′ OH groups on the P1 duplex of the Tetrahymena ribozyme: intrinsic binding energy in an RNA enzyme. Biochemistry *36*, 2465–2477.

Nishimasu, H., Ran, F.A., Hsu, P.D., Konermann, S., Shehata, S.I., Dohmae, N., Ishitani, R., Zhang, F., and Nureki, O. (2014). Crystal structure of Cas9 in complex with guide RNA and target DNA. Cell 156, 935–949.

Ohnishi, Y., Tamura, Y., Yoshida, M., Tokunaga, K., and Hohjoh, H. (2008). Enhancement of allele discrimination by introduction of nucleotide mismatches into siRNA in allele-specific gene silencing by RNAi. PLoS ONE 3, e2248.

Østergaard, M.E., Southwell, A.L., Kordasiewicz, H., Watt, A.T., Skotte, N.H., Doty, C.N., Vaid, K., Villanueva, E.B., Swayze, E.E., Bennett, C.F., et al. (2013). Rational design of antisense oligonucleotides targeting single nucleotide polymorphisms for potent and allele selective suppression of mutant Huntingtin in the CNS. Nucleic Acids Res 41, 9634–9650.

Pattanayak, V., Lin, S., Guilinger, J.P., Ma, E., Doudna, J.A., and Liu, D.R. (2013). High-throughput profiling of off-target DNA cleavage reveals RNA-programmed Cas9 nuclease specificity. Nat Biotechnol 31, 839–843.

Pattanayak, V., Ramirez, C.L., Joung, J.K., and Liu, D.R. (2011). Revealing off-target cleavage specificities of zinc-finger nucleases by in vitro selection. Nat Methods 8, 765–770.

Pedersen, L., Hagedorn, P.H., Lindholm, M.W., and Lindow, M. (2014). A Kinetic Model Explains Why Shorter and Less Affine Enzyme-recruiting Oligonucleotides Can Be More Potent. Mol Ther Nucleic Acids 3, e149.

Pfister, E.L., Kennington, L., Straubhaar, J., Wagh, S., Liu, W., DiFiglia, M., Landwehrmeyer, B., Vonsattel, J.-P., Zamore, P.D., and Aronin, N. (2009). Five siRNAs targeting three SNPs may provide therapy for three-quarters of Huntington's disease patients. Curr Biol 19, 774–778.

Porteus, M. (2016). Genome Editing: A new approach to human therapeutics. Annu. Rev. Pharmacol. Toxicol. 56, 163–190.

Ran, F.A., Hsu, P.D., Lin, C.-Y., Gootenberg, J.S., Konermann, S., Trevino, A.E., Scott, D.A., Inoue, A., Matoba, S., Zhang, Y., et al. (2013). Double nicking by RNA-guided CRISPR Cas9 for enhanced genome editing specificity. Cell 154, 1380–1389.

Ross, P.D., and Sturtevant, J.M. (1960). The kinetics of double helix formation from polyriboadenylic acid and polyribouridylic acid. Proc Natl Acad Sci USA 46, 1360–1365.

Salomon, W.E., Jolly, S.M., Moore, M.J., Zamore, P.D., and Serebrov, V. (2015). Singlemolecule imaging reveals that argonaute reshapes the binding properties of its nucleic acid guides. Cell 162, 84–95.

Schirle, N.T., and MacRae, I.J. (2012). The crystal structure of human Argonaute2. Science 336, 1037–1040.

Schirle, N.T., Sheu-Gruttadauria, J., and MacRae, I.J. (2014). Structural basis for microRNA targeting. Science 346, 608–613.

Schwarz, D.S., Ding, H., Kennington, L., Moore, J.T., Schelter, J., Burchard, J., Linsley, P.S., Aronin, N., Xu, Z., and Zamore, P.D. (2006). Designing siRNA that distinguish between genes that differ by a single nucleotide. PLoS Genet 2, e140.

Sheng, G., Zhao, H., Wang, J., Rao, Y., Tian, W., Swarts, D.C., van der Oost, J., Patel, D.J., and Wang, Y. (2014). Structure-based cleavage mechanism of Thermus thermophilus Argonaute DNA guide strand-mediated DNA target cleavage. Proc Natl Acad Sci USA 111, 652–657.

Slaymaker, I.M., Gao, L., Zetsche, B., Scott, D.A., Yan, W.X., and Zhang, F. (2016). Rationally engineered Cas9 nucleases with improved specificity. Science 351, 84–88.

Sternberg, S.H., LaFrance, B., Kaplan, M., and Doudna, J.A. (2015). Conformational control of DNA target cleavage by CRISPR-Cas9. Nature 527, 110–113.

Sternberg, S.H., Redding, S., Jinek, M., Greene, E.C., and Doudna, J.A. (2014). DNA interrogation by the CRISPR RNA-guided endonuclease Cas9. Nature 507, 62–67.

Tomari, Y., and Zamore, P.D. (2005). Perspective: machines for RNAi. Genes Dev 19, 517–529.

Tomari, Y., Du, T., and Zamore, P.D. (2007). Sorting of Drosophila small silencing RNAs. Cell 130, 299–308.

Tsai, S.Q., Zheng, Z., Nguyen, N.T., Liebers, M., Topkar, V.V., Thapar, V., Wyvekens, N., Khayter, C., Iafrate, A.J., Le, L.P., et al. (2015). GUIDE-seq enables genome-wide profiling of off-target cleavage by CRISPR-Cas nucleases. Nat Biotechnol 33, 187–197.

Ui-Tei, K., Naito, Y., Nishi, K., Juni, A., and Saigo, K. (2008a). Thermodynamic stability and Watson-Crick base pairing in the seed duplex are major determinants of the efficiency of the siRNA-based off-target effect. Nucleic Acids Res 36, 7100–7109.

Ui-Tei, K., Naito, Y., Zenno, S., Nishi, K., Yamato, K., Takahashi, F., Juni, A., and Saigo, K. (2008b). Functional dissection of siRNA sequence by systematic DNA substitution: modified siRNA with a DNA seed arm is a powerful tool for mammalian gene silencing with significantly reduced off-target effect. Nucleic Acids Res 36, 2136–2151.

Vaish, N., Chen, F., Seth, S., Fosnaugh, K., Liu, Y., Adami, R., Brown, T., Chen, Y., Harvie, P., Johns, R., et al. (2011). Improved specificity of gene silencing by siRNAs containing unlocked nucleobase analogs. Nucleic Acids Res 39, 1823–1832.

Wang, X.-H., Aliyari, R., Li, W.-X., Li, H.-W., Kim, K., Carthew, R., Atkinson, P., and Ding, S.-W. (2006). RNA interference directs innate immunity against viruses in adult Drosophila. Science 312, 452–454.

Wang, Y., Juranek, S., Li, H., Sheng, G., Wardle, G.S., Tuschl, T., and Patel, D.J. (2009). Nucleation, propagation and cleavage of target RNAs in Ago silencing complexes. Nature 461, 754–761.

Wee, L.M., Flores-Jasso, C.F., Salomon, W.E., and Zamore, P.D. (2012). Argonaute Divides Its RNA Guide into Domains with Distinct Functions and RNA-Binding Properties. Cell 151, 1055–1067.

Wetmur, J.G., and Davidson, N. (1968). Kinetics of renaturation of DNA. J Mol Biol 31, 349–370.

Wightman, B., Ha, I., and Ruvkun, G. (1993). Posttranscriptional regulation of the heterochronic gene lin-14 by lin-4 mediates temporal pattern formation in C. elegans. Cell 75, 855–862.

Wittrup, A., and Lieberman, J. (2015). Knocking down disease: a progress report on siRNA therapeutics. Nat Rev Genet 16, 543–552.

Wu, X., Kriz, A.J., and Sharp, P.A. (2014). Target specificity of the CRISPR-Cas9 system. Quant Biol *2*, 59–70.

Yin, H., Song, C.-Q., Dorkin, J.R., Zhu, L.J., Li, Y., Wu, Q., Park, A., Yang, J., Suresh, S., Bizhanova, A., et al. (2016). Therapeutic genome editing by combined viral and non-viral delivery of CRISPR system components in vivo. Nat Biotechnol 34, 328–333.

Yu, D., Pendergraff, H., Liu, J., Kordasiewicz, H.B., Cleveland, D.W., Swayze, E.E., Lima, W.F., Crooke, S.T., Prakash, T.P., and Corey, D.R. (2012). Single-stranded RNAs use RNAi to potently and allele-selectively inhibit mutant huntingtin expression. Cell 150, 895–908.

## References

Salomon, W.E., Jolly, S.M., Moore, M.J., Zamore, P.D., and Serebrov, V. (2015). SingleMolecule Imaging Reveals that Argonaute Reshapes the Binding Properties of Its Nucleic Acid Guides. Cell 162, 84–95.

